# ABCH2 transporter in the first line of defense protects malaria vectors from pyrethroids

**DOI:** 10.1101/2023.02.21.529359

**Authors:** Mary Kefi, Vasileia Balabanidou, Chara Sarafoglou, Jason Charamis, Gareth Lycett, Hilary Ranson, Giorgos Gouridis, John Vontas

## Abstract

Contact insecticides are primarily used for the control of *Anopheles* malaria vectors. These chemicals penetrate mosquito legs and other appendages, the first barrier to reaching their neuronal targets. An ATP-Binding Cassette transporter from the H family (ABCH2) is highly expressed in *Anopheles coluzzii* legs, and further induced upon insecticide exposure. RNAi-mediated silencing of the ABCH2 caused a significant increase in deltamethrin mortality compared to control mosquitoes, coincident with a corresponding increase in ^14^C-deltamethrin penetration. RT-qPCR analysis and immunolocalization revealed that ABCH2 is mainly localized in the legs and head appendages, and more specifically, the apical part of appendage epidermis, underneath the cuticle. To unravel the molecular mechanism underlying the role of ABCH2 in modulating pyrethroid toxicity, two hypotheses were investigated: An indirect role, based on the orthology with other insect ABCH transporters involved with lipid transport and deposition of CHC lipids in *Anopheles* legs which may increase cuticle thickness, slowing down the penetration rate of deltamethrin; or the direct pumping of deltamethrin out of the organism. Evaluation of the leg cuticular hydrocarbon (CHC) content showed that this was not affected by ABCH2 silencing, indicating this transporter in is not associated with the transport of leg CHCs. Homology-based modeling suggested that the ABCH2 half-transporter adopts a physiological homodimeric state, in line with its ability to hydrolyze ATP *in vitro* when expressed on its own in insect cells. Docking analysis revealed a deltamethrin pocket on the homodimeric transporter. Furthermore, deltamethrin-induced ATP hydrolysis in ABCH2-expressing cell membranes, further supports that deltamethrin is indeed a ABCH2 substrate. Overall, our findings pinpoint ABCH2 as a key regulator of deltamethrin toxicity.

## Introduction

Malaria is a major impediment to health and prosperity in the Global South. Its prevention is best achieved by vector control which relies heavily on insecticides (1). Malaria incidence halved between 2000-2015, with the majority of the reduction attributed to the use of insecticides (1). However, insecticide resistance is a critical threat to vector control, as some mosquito populations now manifest a striking intensity of the resistance phenotype (2).

Due to the mode of insecticide delivery in vector control, *Anopheles* legs are the key sites for contact insecticide uptake, representing the first barrier to be crossed in the insect (3). Recent studies highlighted the role of mosquito legs in insecticide resistance via structural alterations which reduce the penetration rate of pyrethroids (4) and the overexpression of sensory appendage proteins possibly sequestering the pyrethroid insecticide (5). An ABCH transporter was also identified in mosquito legs through proteomic and transcriptomic analysis (4, 6) and found to be induced by a short-term deltamethrin exposure in *An. coluzzii* (6).

ABC transporters are present in all kingdoms of life functioning as primary-active transporters energized by ATP hydrolysis (7) in the highly conserved nucleotide binding domains (NBDs) that are associated with the translocator (8) transmembrane domains (TMDs) (8). ABC transporter involvement in insecticide resistance and toxicity has been suggested in some studies (6, 9–12), and at the same time a differential expression of subsets of ABC proteins in pyrethroid-resistant *Anopheles* mosquito populations has also been reported (13, 14). Additionally, a number of investigations support the up-regulation of ABC transporters from pests, such as *B. tabaci* and *P. xylostella* after exposure to different classes of insecticides, implicating a correlation between pesticide detoxification and transport (15, 16). ABC transporters from the C-subfamily have been implicated in pest resistance to insecticidal pore-forming proteins from *Bacillus thuringiensis* (Bt). Recently, a multidrug resistance-associated ABC, *Cp*MRP of the polar leaf beetle *Chrysomela populi*, was identified as the first candidate involved in the sequestration of phytochemicals in insects (17, 18). An RNAi toxicology screen in *Drosophila* implicated a C family ABC transporter (CG4562) in spinosad transport, as well as pinpointing the role of the P-glycoprotein orthologue *Mdr*65 as the most important ABC in terms of chemoprotection (19). Much of this functional analysis of the interaction of ABC transporters with insecticides has been studied through heterologous expression in insect cells and *Xenopus* oocytes (20, 21).

Transporters of the H sub-family are present in all insects and other arthropods (22), *Dictyostelium* and zebrafish, but are absent from plants, worms, yeast, or mammalian genomes (23). The H sub-family transporter members are export proteins sharing similarities with members of the G group (8). They are composed of a single NBD and TBD, hence they are half-transporters, which means they need to dimerize to be functional (24, 25). In *An. coluzzii*, three genes encode for ABCH transporters (*ABCH1, ABCH2* and *ABCH3*) (25). ABCH transporters have been implicated in lipid transport, as knock-down experiments in *Drosophila melanogaster, Tribolium castaneum, Locusta migratoria, Plutella xylostella and Nezzara viridula* result in high lethality due to desiccation (26–30). Immunofluorescence analysis showed that the *Drosophila Snu* protein, an ABCH2 orthologue, localizes to the apical plasma membrane of the epidermal cells of larvae. ABCH transporters have been found differentially expressed in insecticide resistant populations or after insecticide exposure (6, 10, 31–35), however whether their implication in such phenomena is linked to the direct binding/transport of insecticides or a consequence of other physiological roles remains elusive.

The *An. coluzzii* ATP-Binding-Cassette Transporter, *ABCH2* (*ACON003680*), an ABC transporter of the H family was the only ABC transporter identified to be expressed in mosquito legs in both proteomic and transcriptomic datasets (4, 6). Overexpression of this transporter in the legs, upon a short-term exposure of *An. coluzz*ii mosquitoes to deltamethrin (6) provided evidence to implicate this transporter in pyrethroid toxicity.

Here, we aimed to elucidate the basis of the resistance conferred by the *An. coluzzii* ABCH2 transporter to deltamethrin toxicity.

## Results

### RNAi-mediated silencing of *ABCH2* revealed its implication in pyrethroid toxicity

Former work of our group showed that *ABCH2* was induced upon deltamethrin exposure in mosquito legs (6). ABCH2 was previously identified uniquely in the leg proteome of *An. coluzzii* (4) and as well in the leg transcriptome (6). Here, the expression in different tissues, both at the transcript and protein levels was determined. The transcript abundance in different dissected body parts and tissues of 3-5 day-old female *An. coluzzii* was evaluated with RT-qPCR, with the expression of all samples being normalized against the expression of abdominal walls. As expected, the *ABCH2* relative expression in legs is greater than other dissected tissues (Figure S1A). The protein abundance of ABCH2 is higher in the legs with significant protein detected in heads too, as evidenced by western blot analysis (Figure S1B). The ABCH2 expression in head is detected to specific head appendages (antennae, proboscis, maxillary palps) (Figure S1C) and not to the rest head.

To test whether *ABCH2* exhibits phenotypes related to pyrethroid toxicity, we performed RNAi-mediated silencing in newly emerged adults of an insecticide resistant strain coupled to bioassays. dsRNA specifically targeting *ABCH2* transcripts were designed, generated and introduced intrathoracically into newly emerged *An. coluzzii* females via nano-injections. The silencing efficiency was evaluated both at the transcript and protein levels. According to RT-qPCR, dsRNA-mediated silencing reduced *ABCH2* transcript levels in whole female mosquitoes by approximately 75% (Figure 1A). Western blot analysis using the ABCH2 peptide antibody detected a specific signal at approximately 85 kDa, the expected size of the transporter and was used to prove the silencing efficiency at the protein level. Indeed, ABCH2 protein in legs and head appendages (proboscis, antennae, and maxillary palps) of female mosquitoes were barely detectable compared to ds*GFP* counterparts (Figure 1B). The effects of *ABCH2* silencing in pyrethroid toxicity were evaluated via deltamethrin toxicity assays. Mosquito knock-down 1 hour post exposure in ds*GFP* injected mosquitoes was 23% while in dsA*BCH2* mosquitoes, it was 89.7%, showing a significant increase (Student’s t-test, *p*-value<0.0001). The ds*ABCH2* mosquitoes did not recover, with a mortality of 98% after 24 hours, while in ds*GFP* controls, the mortality was 55% (Figure 1C, D).

**Figure 1.**
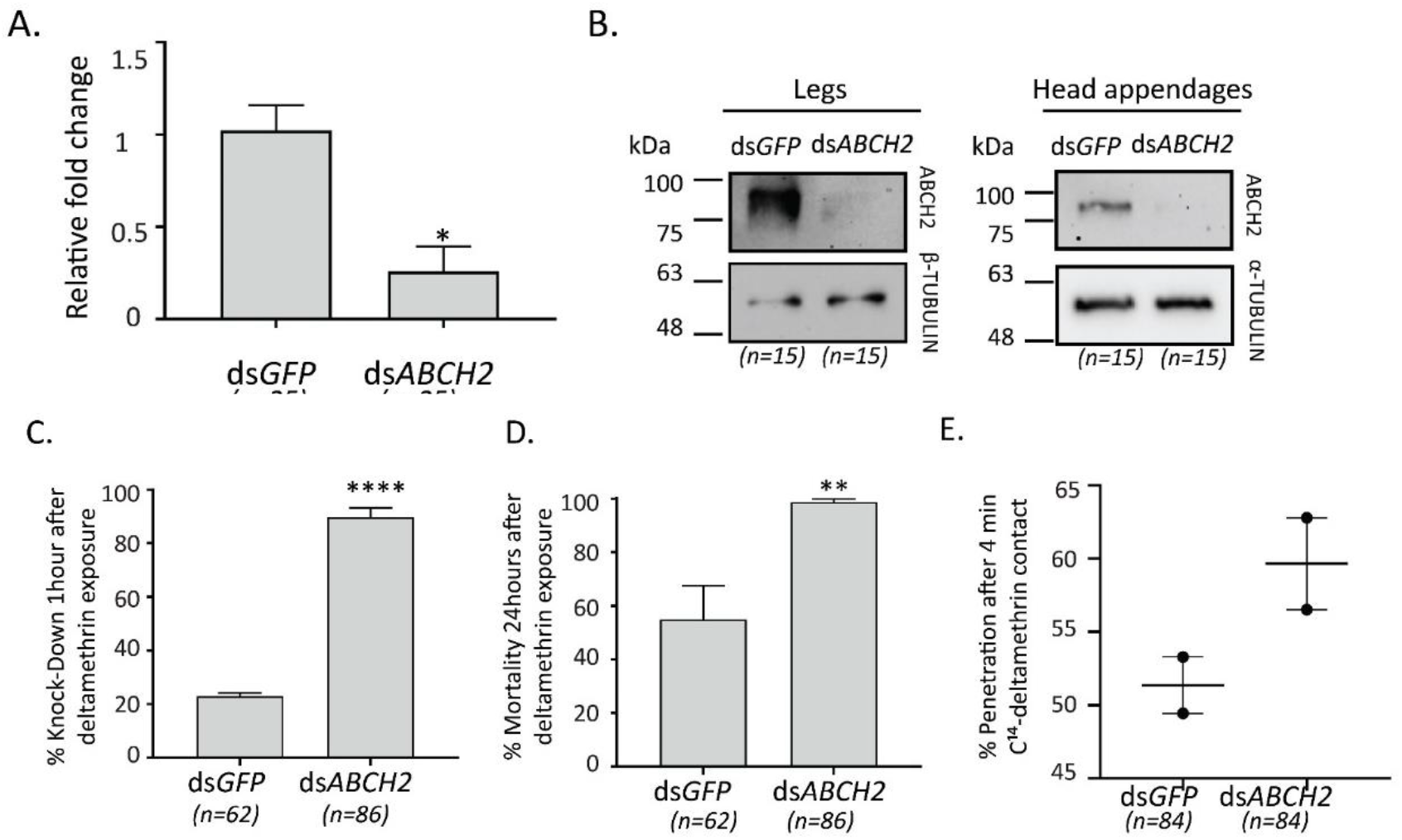
RNAi silencing efficiency and deltamethrin toxicity assays. A. Silencing efficiency estimation of ds*ABCH2* against ds*GFP* injected females with RT-qPCR. ABCH2 transcript level reduction by 74,7% accomplished, as indicated by the mean of 3 biological replicates +SEM; *P*-value = 0,0114 (*) determined by t-test, B. Western Blot analysis of leg and head appendages protein extracts 3 days post injections, verified the reduced protein levels of ds*ABCH2* against ds*GFP* injected controls. Alpha or beta-TUBULIN were used as loading control, C. % Knock down of ds*ABCH2* and ds*GFP* female *An. coluzzii*, subjected, 72h post injection, to 1 hour deltamethrin exposure (0.016%); Mean_(ds*GFP*)_ =23% + 1.33 and Mean_(ds*ABCH2*)_ =89.6% + 3.66 for n= 4 biological replicates; *P*-value<0.0001 (****), D. % mortality 24 hours post exposure, Mean_(ds*GFP*)_ = 55% +12.58 and Mean_(ds*ABCH2*)_ = 98.75% +1.25, for n= 4 biological replicates; *P*-value= 0.0092 (**), E. % Penetration of C^14^-deltamethrin after 4 minutes of contact in dsGFP and dsABCH2 mosquitoes. Penetration corresponds to the ratio of the internal to the total counts in this time point. Mean of two biological replicates (n=84 mosquitoes/condition).

To rationalize the increased deltamethrin mortality, we determined the amount of internalized deltamethrin by using ^14^ C-deltamethrin contact toxicity assays on silenced and control *An. coluzzii*. dsABCH2 mosquitoes exhibit increased ^14^ C-deltamethrin penetration compared to their dsGFP counterparts by about 10% in the two biological replicates (Figure 1E).

### ABCH2 is localized on the leg/appendage epidermis, underneath the cuticle, with apical polarity

As ABCH2 expressed in leg and head appendages is implicated in deltamethrin toxicity, our ensuing objective was to determine its (sub)cellular localization in such tissues to gain more insights regarding its role in insecticide toxicity. Towards this, the specific antibody against this transporter was used in immunofluorescence experiments in leg cryo-sections. We repeatedly obtained a specific signal (green) underneath the cuticle as shown in the merged-fluorescent images with the bright-field channel (Figure 2A, B right bottom). The signal observed followed the linear contour of the leg, just underneath the cuticle and adjacent to the nuclei stained red with TO-PRO-3 dye. Also stained were characteristic triangle-shaped protrusions towards the cuticle. To confirm this location as sub-cuticular epithelia, we obtained a similar staining pattern on leg cryo-sections with an E-cadherin antibody, known to stain these cells (36–39) (Figure 2C,D). The ABCH2 signal appeared to be located apically, as opposed to the more even distribution of Cadherin over the whole plasma membrane.

**Figure 2.**
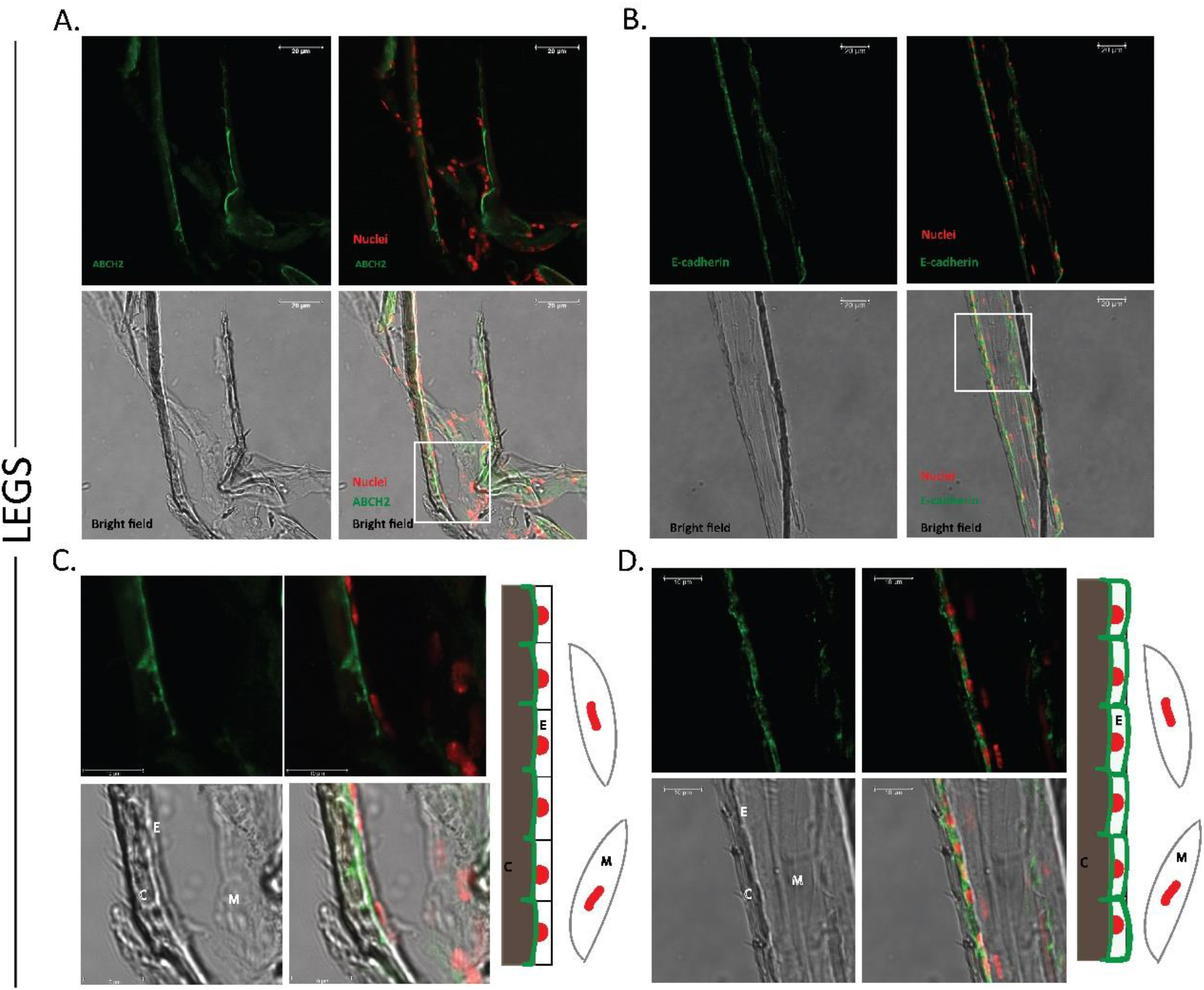
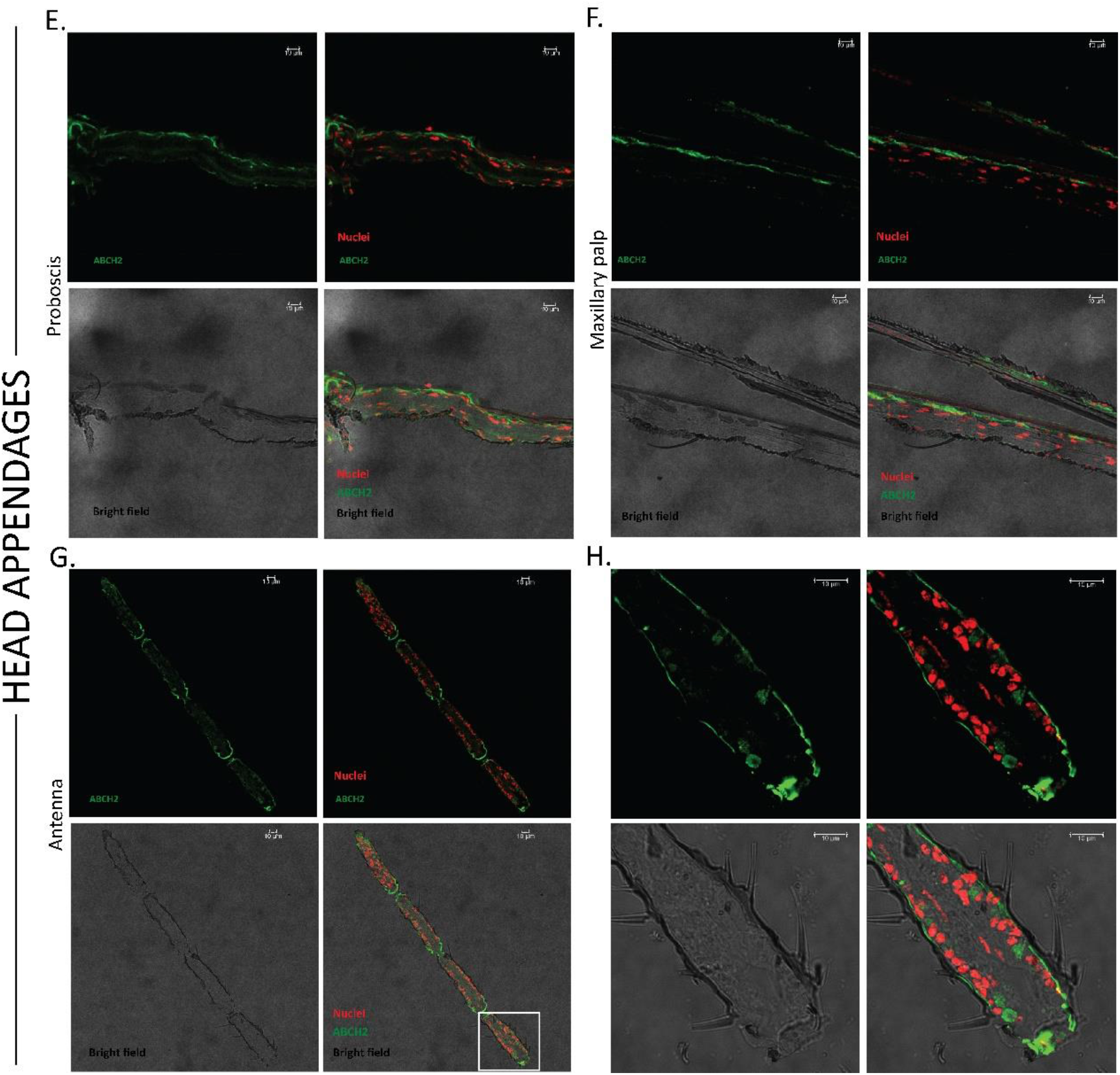
Sub-cellular localization of ABCH2 transporter. Immunohistochemical images from longitudinal leg cryosections of 3-5 day old female *An. coluzzii*. A. ABCH2 localization on epidermal cells, underneath the cuticle, polarized towards the apical side. B. Epithelial staining on leg cryosections with a marker-antibody against E-cadherin validating the presence of an epidermal layer underneath the cuticle. C, D. Zoomed images of the selected, white squares of figures A and B respectively, together with graphical depictions of the main structures observed. E,F,G. ABCH2 localization on epidermal cells of head appendages: proboscis, maxillary palp and antenna respectively. H. Zoomed image of the selected, white square of figure G. Images were obtained with confocal microscope (40x). RED: Cell nuclei are stained with TO-PRO-3; Green: antibody staining; Merged images with and without bright-field channel are also depicted; C=cuticle, E=epidermis, M= muscles; scale bars of 10-20 μm are illustrated.

A similar ABCH2 staining pattern was also observed in cryo-sections of head appendages (Figure 2E,F,G,H). Furthermore, using the DeepLoc-1.0 tool (40), we predicted a plasma membrane sub-cellular localization for this multi-span transmembrane protein (Figure S2). These lines of evidence support that mosquito ABCH2 is localized on leg/appendage epidermal cells, most probably apically polarized towards the cuticular structure.

### Deltamethrin toxicity is not attributed to CHC differences in the legs

As several studies implicate ABCH transporters in lipid transport, we next sought to test whether the increased mortality and penetration in ABCH2 depleted mosquitoes is due to reduction of lipid species. *An. coluzzii, D. melanogaster, L. migratoria, P. xylostells* and *T. castaneum* possess three *ABCHs* (with one-to-one orthology relationships. Interestingly, all ABCHs with reported roles in lipid transport are clustered within the same clade with AcABCH2 (Figure S3). Thus, we wondered whether ABCH2 also participates in transport of cuticular hydrocarbons (CHCs), being the most abundant lipid species in *Anopheles* leg cuticle (41) and having a documented role in reducing the penetration rate of insecticides (41). To address this, we performed CHC analysis in mosquito legs from control or dsABCH2 injected mosquitoes. The analysis of hexane leg extracts indicated no significant difference either in the total CHC content between ds*ABCH2* and ds*GFP* mosquito legs (Figure 3), or in the relative abundance of individual CHC species (Figure S4). These results suggest that the increase in deltamethrin toxicity observed in the ABCH2-silenced mosquitoes is not due to a change in the amount or profile of epicuticular hydrocarbons.

**Figure 3.**
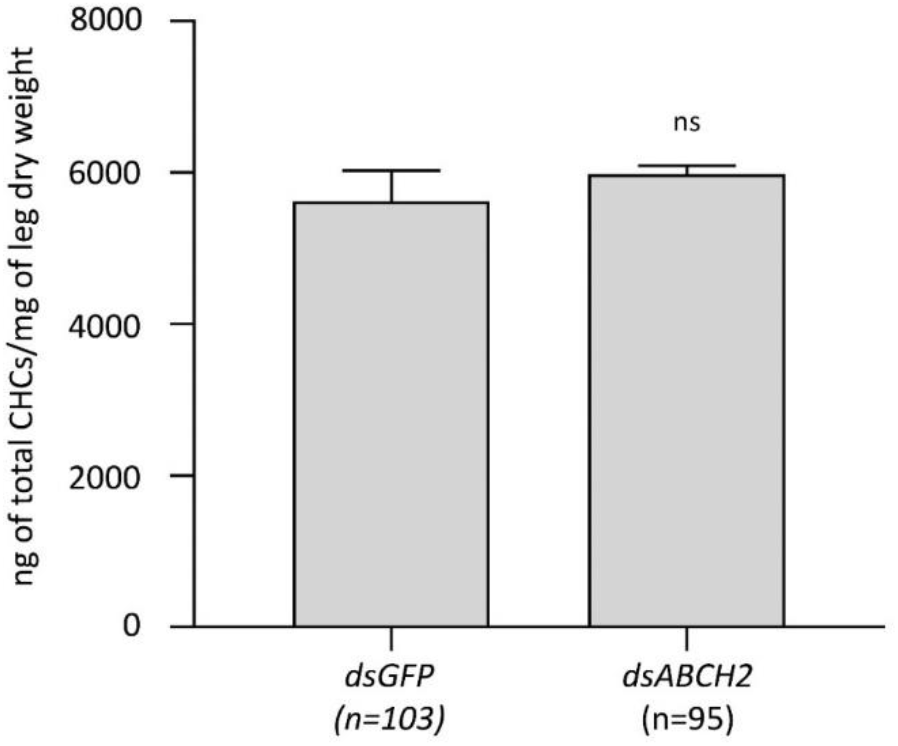
Total Cuticular Hydrocarbon content of legs derived from ds*ABCH2* and ds*GFP* injected mosquitoes. Average total CHC content from three replicates analyzed for each condition (95 ds*ABCH2* and 103 ds*GFP* female mosquitoes, 30-35 mosquitoes per replicate); Mean + SEM; Mean_(ds*GFP*)_ =5626.33+ 401.5 and Mean_(ds*ABCH2*)_ =5981.67+ 112.5 for n= 3 biological replicates; *P*-value=0.4422 (non-significant, ns).

### *In silico* and *in vitro* tools provide evidence for a membrane-bound ABC transporter which functions most probably as a homodimer

The ABCH2 sequence was modelled after the homodimeric ABCG1 structure (42), identified to be its closest homologue with an available structure. This consented to identify contacts stabilizing the inter-protomer interfaces using the Protein Interaction Calculator (PIC) web-server (43). The number and nature of interactions stabilizing the dimeric ABCG1 interface is equivalent to those occurring in the modelled ABCH2 one (Figure 5A and 5B and Supplementary File 2), suggesting that ABCH2 adopts a homodimeric state. To verify this, we expressed it using the baculovirus-mediated system and assessed its activity *in vitro*. After validating the specific expression in ABCH2-infected cells compared to cells infected with an empty-bacmid (Figure S5A), we proceeded to sub-cellular fractionation. As expected (Figure S5A,B and Figure 4C), the transporter is only present in the membrane fractions of the ABCH2-infected cells. Inverted Membrane Vesicles (IMVs) isolated from such cells, indicate an ATP hydrolysis specific to ABCH2, as the increase of Pi released was statistically significant compared to IMVs free of ABCH2, expressing an unrelated membrane protein (Figure 4D, Figure S6). In order to approximate the rate of ABCH2-related ATP hydrolysis, we performed western blot analysis relying on measured quantities of a purified His-tagged protein, used as a reference to estimate the amount of His-ABCH2 in expressing IMVs (Figure S7). Based on this estimate, free ABCH2 hydrolyzes ~178pmoles Pi/pmol ABCH2/minute (Table S2), representing a substantial basal hydrolysis rate compared to other membrane-embedded motor proteins (44, 45).

**Figure 4.**
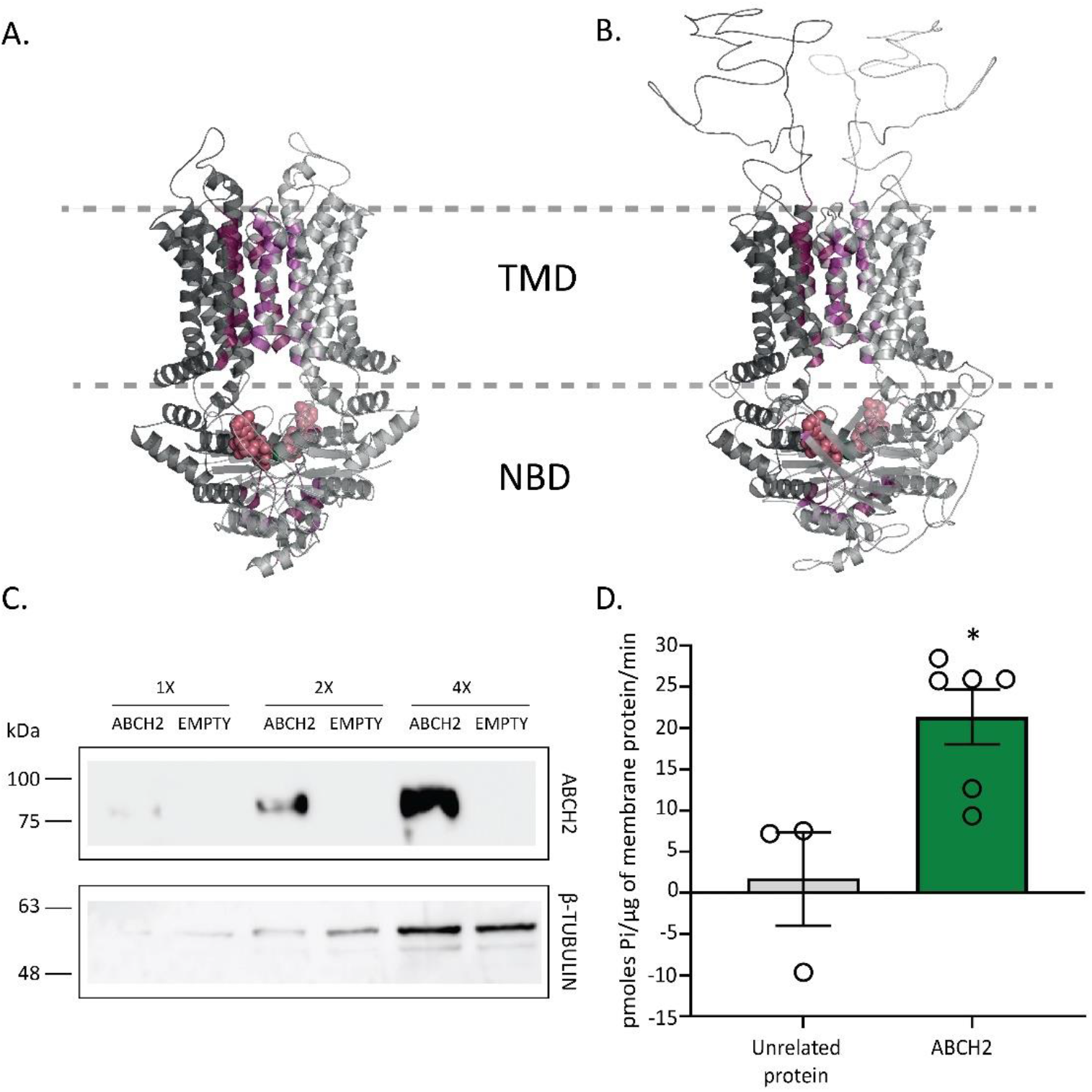
Functional expression of ABCH2, as a homodimer. A and B. Interface residues participating in interactions stabilizing the homo-dimeric interface are presented with purple surface colors. Bound ATPs in the Nucleotide Binding Domains (NBDs) are presented with red spheres. The two protomers in the ABCG1 structure (A) or in the modelled ABCH2 (B) are distinguished by two grey scales. C. Western blot analysis using ABCH2 antibody in whole membrane preparations of ABCH2- and Empty Bacmid-expressing *Sf9* cells. Different concentrations (1x, 2x, 4x) of membrane preparations were tested with B-TUBULIN serving as loading control. D. ATPase activity of membrane preparations using malachite green. Mean + SEM of six biological replicates in ABCH2-membranes and 3 biological replicates in unrelated protein-expressing membranes and the average of 2-3 technical replicates for each biological is presented in bars. Mean + SEM; Mean_(EMPTY)_ =1.7+ 5.67 and Mean_(ABCH2)_ =21.37+ 3.32; *P*-value=0.0148 (*).

**Figure 5.**
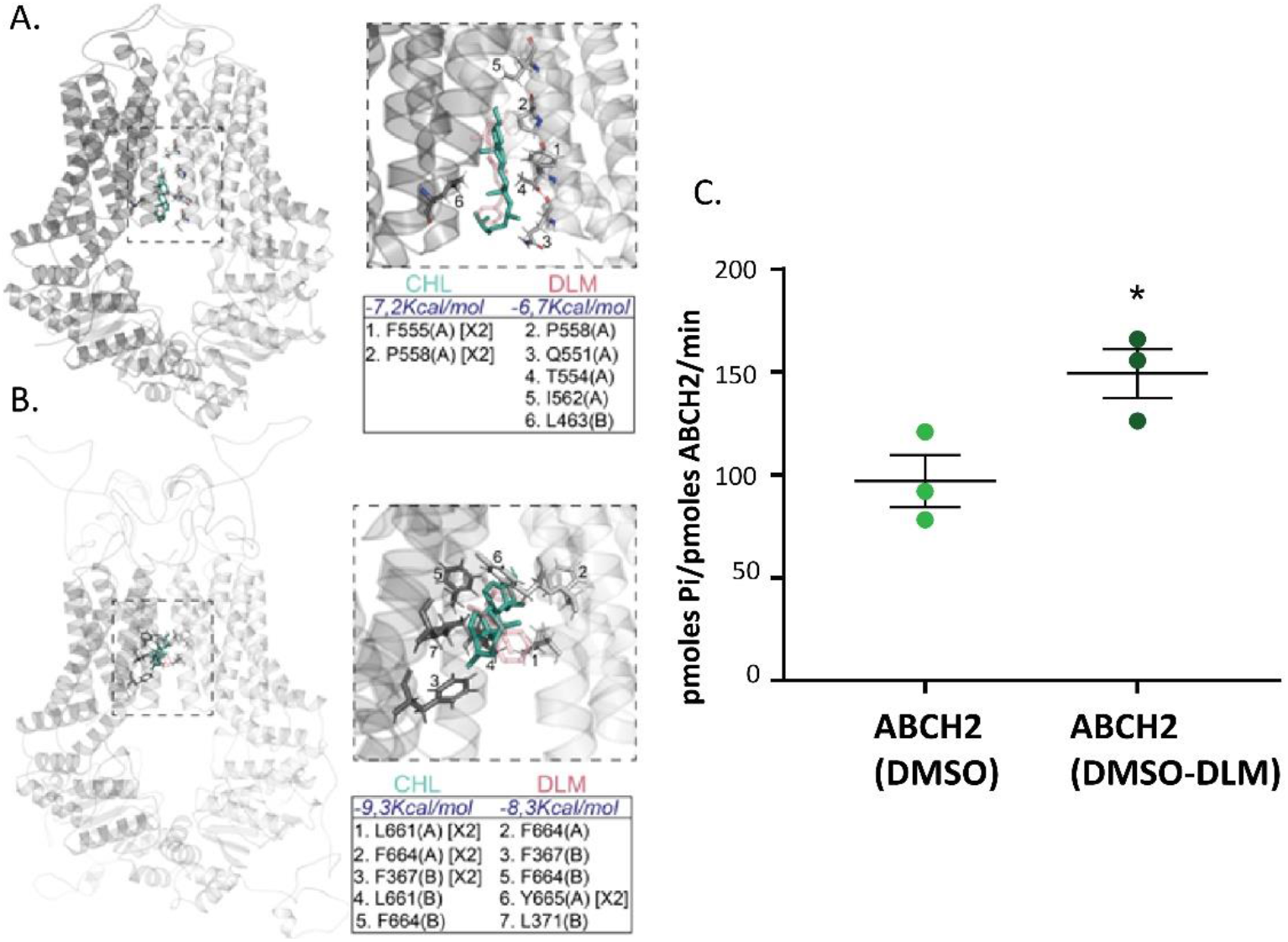
A. ABCG1 having docked cholesterol (CHL) and deltamethrin (DLM) as described in the text and methods. CHL is presented with green sticks, whereas DHL with transparent red sticks. Zoom-in of the indicated dotted area is presented (right top panel) together with the calculated energies and substrate-stabilizing interactions (right bottom panel). The residues participating in the hydrophobic interactions with the substrates are given in the table for each protomer (A) or B), and the number of hydrophobic bonds for each residue is indicated in brackets (when more than one), B. As in A with the modelled ABCH2 structure, C. ATPase activity of membrane preparations using deltamethrin. Pmoles of Pi produced per pmole of ABCH2 per minute of reaction are depicted in ABCH2-expressing IMVs in reaction containing the organic solvent (DMSO) and deltamethrin (DMSO-DLM). Mean +SEM of three biological replicates and the average of 2 technical replicates for each biological are depicted. Mean_(DMSO)_ =97.07+12.5 and Mean_(DMSO-DLM)_ =149.3 + 11.9; *P*-value=0.0394 (*).

### Docking and *in vitro* experiments indicate that deltamethrin is a putative substrate of ABCH2

To validate whether deltamethrin can act as an ABCH2 substrate, we initially performed docking analysis *in silico*. The model used ABCG1 has been crystallized in many distinct cholesterol-liganded states (42). In the nucleotide-free inward facing conformation (PDB:7R8D), cholesterol molecules have been allocated to the transmembrane part of the transporter. Two of them at the channel interior and three additional ones at its exterior. In the ATP bound state (PDB:7R8E), the cholesterol molecules were observed uniquely in the channel exterior. We assessed the cholesterol binding pockets via the Protein-Ligand Interaction Profiler Web-server and focused our subsequent analysis on the most well defined, interior pockets, present within the translocation path (42). Initial protein-ligand docking experiments were carried out using AutoDock Smina (46) to retrieve the binding energies of cholesterol and deltamethrin. From the top scores, we present (Figure 4A) the ones having binding modes resembling closely the crystal structure (42). The results indicate that both ligands docked to the pocket with similar energies (Figure 5A). Moreover, the number and nature of interaction stabilizing the two ligands into the pocket is equivalent (Figure 5A). These observations suggest deltamethrin acting as a potential substrate of ABCG1.

Next, to assess whether deltamethrin is an ABCH2 substrate, we performed protein-ligand docking experiments on the modeled ABCH2 without a bias related to the substrate pocket. Thus, we widened our search grid to include the entire translocator (i.e. transmembrane region). The top scores that emerged indicate a substrate pocket in the translocator interior, just above the one defined by the crystal structure (compare Figure 5A *vs* B, left panels). In such a pocket, both cholesterol and deltamethrin dock with equivalent orientation, energies and interactions (Figure 5B).

To look for experimental evidence of binding, deltamethrin was introduced in the *in vitro* assay to ask whether it stimulates the basal ABCH2-IMVs ATPase activity. Interestingly, a significant stimulation over the basal activity was observed when deltamethrin was included in the reaction (Figure 5C). Taking all together, we propose deltamethrin to be a physiologically relevant ABCH2 substrate.

## Discussion

Mosquito legs/appendages are the first line of defense against insecticides when *Anopheles* vectors come in contact with impregnated bed-nets and wall surfaces (47, 48). The presence of transport systems in appendages apart from serving key physiological functions, could be involved in conferring tolerance against chemicals, such as insecticides (14). But whether and how mosquito transporters are implicated in drug toxicity remains largely elusive. Here, we show that the ABCH2 transporter has an essential role in alleviating pyrethroid toxicity in *An. colluzii*, presumably by direct transport of pyrethroids out of cuticular epithelial cells, based on several lines of evidence.

Firstly, RNAi-mediated silencing of the *ABCH2*, an ABC transporter previously shown to be induced by deltamethrin, increased pyrethroid toxicity substantially (from 23% to 90% knocked-down mosquitoes). Moreover, ^14^ C-Deltamethrin penetration experiments revealed that *An. coluzzii* lacking the transporter showed increased penetration of radiolabeled insecticide after short (4 min) contact. It is plausible that the slower insecticide penetration rate, provided by pumping out the insecticide, gives greater time for metabolic enzymes to detoxify the insecticide, making the transporter-devoid mosquitoes far more vulnerable to insecticides.

Secondly, the anatomical localization, in relevance to insecticide uptake by legs and appendages and the cellular localization within epidermal cells with an apical polarity, oriented towards the cuticle (39), is consistent with the ability of the transporter to remove insecticides from the organism at the first point of entry. The ABCH2 orthologue in *D. melanogaster* is also localized in epidermal cells, indicated by epidermal-Gal4/RNAi screening and by localization experiments with fused GFP, showing its topology on the apical surface of larval epidermal cells (49). It is common for transporters to compartmentalize on the membranes towards the site they exhibit their function (50). A polarity towards the outermost part, i.e. the exoskeleton could suggest the ability of the transporter to facilitate transport of cuticular components and/or other substrates, such as insecticides, out of the organism and both scenarios were explored.

Thirdly, we tested the scenario of transport of deltamethrin across cell membranes that would lend support to the difference in penetration in being attributed to direct export from epidermal cells and sequestration of the insecticide into the cuticular structure. Several reports have shown that ABC transporters can participate in phase 0 detoxification (51). To explore this possibility, we performed *in silico* analysis and functional ABCH2 expression *in vitro* which revealed that ABCH2 is a functional transporter of homodimeric nature. ABCHs are half-transporters, meaning they need to form either homo- or hetero-dimers in order to be functional (52). *In silico* data support the homodimerization scenario due to the similar nature and number of interface interactions observed in the modelled ABCH2, as in the homodimeric ABCG1 crystal structure (Figure 4A and B, supplementary file 2). Additionally, the elevated ATP hydrolysis observed in the ABCH2-IMVs in the absence of a substrate (Figure 4C), is indicative of an active transporter with significant basal activity, as previously reported for other ABC transporters (45).

Membrane preparations from recombinant *Baculovirus* infected Sf9 cells have been widely used to detect interactions of compounds with ABC transporters of several families (53–55), however to our knowledge this is the first *in vitro* functional expression of an ABC transporter of the H sub-family. Moreover, evidence of insect ABCH2 orthologues also supports the homodimer role in resistance, since in RNAi screens of [*D. melanogaster* (49, 56), *N. viridula* (29), *T. castaneum* (26)] mortality was observed only with ABCH2-orthologues and no other paralogues exhibited similar phenotypes (mortality),.

Our ensuing objective was to test whether deltamethrin is an ABCH2 substrate. In general, providing direct proof of the ability of a transporter to translocate a specific molecule is challenging (57). Decades of research in biological transport systems have highlighted the difficulty in assigning a substrate to an ABC transporter and the low reproducibility arising when using different systems (57). As a first indication that deltamethrin could be a substrate of ABCH2, we performed protein-ligand docking analysis on the modeled ABCH2 structure and demonstrated that, deltamethrin could act as an ABCH2 ligand by the identification of a putative binding pocket. To further substantiate that deltamethrin is indeed an ABCH2 substrate, we performed ATP hydrolysis measurements on isolated membrane vesicles and we observed a substantial stimulation over its basal activity in the presence of deltamethrin. Based on *in silico* and *in vitro* evidence, we thus tentatively conclude that deltamethrin is indeed an ABCH2 substrate.

Lastly, we explored the possibility that the reduced insecticide penetration could be associated with the amount of cuticular lipids on ABCH2-silenced mosquitoes. Based on our phylogenetic analysis, *An. coluzzii* ABCH2 is an orthologue of other insect ABCH transporters implicated in transport of cuticular components in early developmental stages (29). In the insect, knockdown studies have associated ABCH transporters with cuticular lipid transport abnormalities primarily through lipid staining of silenced individuals. Albeit, to our knowledge, the exact molecules transported by this essential protein have not yet been identified (27, 30, 49, 58). Recent work on another ABCH transporter oksyddad (orthologue to ABCH1, Figure S4) indicates that it is required for CHC deposition at the surface of the wing cuticle either directly or indirectly, while it is not implicated in the *D. melanogaster* cuticular barrier formation, shown to be mediated by the ABCH2-orhologue Snu. (56). Another study revealed that the leg epicuticle of insecticide resistant *Anopheles* mosquitoes, which uptake deltamethrin slower upon tarsal contact, is thicker mainly due to enhanced deposition of cuticular hydrocarbons, the major lipid species in leg epicuticles (4). When we analyzed *ABCH2*KD legs versus controls, however, the total hydrocarbon content in both cases were similar and the relative abundances of each CHC species identified also showed non-significant differences. Therefore, the increase in deltamethrin toxicity demonstrated in the *ABCH2*-silenced mosquitoes is not attributed to the reduction of CHCs, at least in the time frame we tested. However, potential transport of CHC species by ABCH2 could happen either earlier in development or at different time points during adult life. Alternatively, we cannot exclude the possibility that ABCH2 facilitates export of other lipid species present in leg epicuticles, such as fatty acids, esters, alcohols, sterols, phenols or other components of the envelope such as wax and cement of unknown composition (59).

Summarizing, several lines of evidence presented here support that deltamethrin could be an ABCH2 ligand (and a putative substrate for translocation): 1) the significant increase in mortality and insecticide penetration in early time points in ABCH2-silenced *An. coluzzii*, 2) the localization of the transporter on the apical membrane of the epidermal layer is in accordance with the hypothesis that this molecule is able to pump out compounds into the exoskeleton and perhaps out of the organism, 3) the *in silico* docking experiments on the modelled ABCH2, derived by the available crystal structure of the ABCG1 homolog, indicates that deltamethrin is a potential ABCH2 substrate, 4) the increased deltamethrin-induced ATP hydrolysis in the *in vitro* system. Based on the above, we propose that ABCH2 mediates deltamethrin efflux in *An. colluzzi*.

Altogether, our results provide evidence for an ABC transporter-based, fast-acting resistance mechanism present at the appendages. As mosquito legs/appendages are a highly epidemiologically relevant tissue with vector control relying on insecticide penetration through the legs, the association of transporter proteins with pyrethroid toxicity and the understanding of the underlying molecular mechanisms would be very useful in vector control innovations. Due to the druggable characteristics of the ABCH2 molecule, specifically, which is an arthropod-specific, plasma membrane-bound transporter present in the epidermis of mosquito appendages, we consider this as a potential target for insecticide formulation add-ons to synergise pyrethroid toxicity.

## Materials and Methods

### Mosquito strains

The mosquito strain used in this study belong to the *An. coluzzii* species complex and was maintained in the laboratory under the same conditions for several generations before analysis. The standard insectary conditions for all strains were 27 °C and 70–80% humidity under a 12-h: 12-h photoperiod with a 1-h dawn:dusk cycle. The strain is derived from Burkina Faso (VK7) and the colony used for the experiment is the VK7-LR colony, which has lost part of its resistance, as described in (6).

### Antibodies

Rabbit polyclonal antibodies targeting *An. coluzzii* ABCH2 peptide (residues 466-482:VKEYYSDLDSALGAVRD) were synthesized and affinity purified by Davids Biotechnologie.E-cadherin mouse antibody was kindly provided by Dr. Siden-Kiamos (60). Mouse anti-beta-tubulin and mouse anti-alpha-tubulin used for normalization control in western blot analysis were purchased by Santa Cruz Biotechnology and Developmental Studies Hybridoma Bank respectively.

### Western Blot Analysis

Dissected tissues from 10-15 female mosquitoes were homogenized into 100-150 μl RIPA Buffer (50Mm Tris pH=8, 150mM NaCl, 1% SDS, 1% NP-40, 1mM EDTA, 1mM EGTA and 1 mM PMSF). After a 10 min centrifugation step at 10,000g the supernatants were immediately supplemented with equal volume (100-150μl) of Laemmli Sample Buffer. Polypeptides were resolved by 12% acrylamide SDS-PAGE and electro-transferred on nitrocellulose membrane (GE Healthcare Whatman). The membranes were subsequently probed with anti-ABCH2 antibody at 1:250 dilution or anti-β-tubulin (Cell Signaling) at 1:500 dilution in a 1% skimmed milk/TBS-0.1%Tween buffer. Antibody binding was detected using goat anti-rabbit or anti-mouse IgG coupled to horseradish peroxidase (Cell Signaling) (dilution: 1:5000 in 1% skimmed milk in TBS-Tween). Visualization was performed using a horseradish peroxidase sensitive ECL western blotting detection kit (SuperSignal West Pico PLUS Chemiluminescent Substrate, ThermoScientific) and the result was recorded using Chemidoc Imaging System (Bio-Rad Laboratories).

### Tissue dissection and relative expression estimation

To determine *ABCH2* expression levels in different tissues, 3-4 day-old female *An. coluzzii* were dissected. Heads, thoraces, guts with the attached malpighian tubules and ovaries, abdominal walls and legs were separated into TRIzol™ Reagent. 20 tissues were placed in 200 μl reagent in three replicates and then manufacturer’s instructions were followed. After elution, RNAs were subjected to DNase I treatment (ThermoFisher Scientific, DNase I, RNase-free). The quantity of DNase-treated RNAs was calculated using a Nanodrop spectrophotometer and equal quantities (1μg) from each sample were used for cDNA synthesis. This was carried out using EnzyQuest Reverse Transcriptase (company) and oligodTs according to the guidelines and the reaction products were used for RT-qPCR. RT-qPCR was performed in a BIO-RAD cycler in the following conditions: 3 min at 95°C, 40 cycles of 15 seconds at 95°C and 30 seconds at 60°C and the analysis included three biological and two technical replicates within each biological replicate. Relative expression was normalized with *An. coluzzii Ribosomal Protein S7* (*S7*) housekeeping gene. The relative expression of *ABCH2* among different tissues was estimated against abdominal wall samples. Graphs were produced and statistically analyzed with GraphPad Prism software version 8 using Student’s t-test.

### Cryosectioning, immunofluorescence and confocal microscopy

Legs and head appendages from 3-5 day-old *An. coluzzii* were dissected and placed for 2 hours in PCR tubes containing 4% PFA in 1x PBS buffer for fixation. After removal of fixative, legs were incubated overnight with 30% sucrose/PBS buffer at 4°C. Legs were then immobilized in eppendorf lids covered with Optimal Cutting Temperature compound (O.C.T - Tissue-Tek SAKURA) and placed at −80°C. 5 μm longitudinal leg sections were obtained in cryostat (Leica CM1850UV) and were placed on superfrost microscope slides (Thermo Scientific). The slides were washed (3 × 5 min) with 0.02% Tween/PBS, followed by a 10 minute incubation with 0.03% Triton/PBS. After 1 hour blocking with 1% Fetal Bovine Serum in 0.03% Triton/PBS, the slides were incubated overnight with the ABCH2 antibody in 1:500 dilution, at 4°C. The next day goat anti-rabbit or goat anti-mouse (Alexa Fluor 488, Molecular Probes) were used in 1:1000 dilutions. TO-PRO-3 dye (Molecular Probes) was used for nuclei staining after RNAse A (Invitrogen™ Ambion™) treatment in a 1:1000 dilution for 5 min. Observation and image attainment were carried out at a Leica SP8 laser-scanning microscope, using a 40x-objective.

### dsRNA design, generation, nano-injections and silencing efficiency

Primers for dsRNA synthesis of *ABCH2* were designed using PrimerBlast and they amplify a product of 525 bp (primer sequence, Table S1). The T7 promoter sequence (5’ TAATACGACTCACTATAGGG 3’) was added at the 5’ end of both forward and reverse oligos. VK7 cDNA was used as template for PCR amplification using Phusion High-Fidelity DNA Polymerase (New England Biolabs), following manufacturer’s instructions. Specific amplification was verified on a 1.5% agarose gel and the rest of the reaction was purified using Macherey-Nagel™ Nucleospin™ Gel and PCR Clean-up Kit. The purified product was used as template for dsRNA synthesis using HiScribe™ T7 High Yield RNA Synthesis Kit (New England Biolabs) with subsequent purification with MEGAclear™ Transcription Clean-Up Kit (Ambion). The purified dsRNAs were diluted to a 3 μg/μl concentration and 69 nl of this were injected into CO_2_-anaesthetized female 0-day mosquitoes. The injections were performed intrathoracically using Nanoinject II Auto-Nanoliter Injector (Drummond Scientific Company). As control a 500bp dsRNA prepared form the *green fluorescent protein* (*GFP*) non-endogenous gene was used after similar preparation (primer sequence, Table S1). Injected mosquitoes were placed in cups and kept in insectary conditions with 10% impregnated sugar cottons for 72 hours. After that their legs were dissected, RNA extracted, cDNA synthesized and RT-qPCR was conducted, as described in detail above. Primers for qPCR were designed out of the region-targeted by dsRNA resulting in 150-200 bp product (primer sequence, Table S1) and according to standard curve construction their efficiency was 96%. Silencing efficiency for each dsRNA was estimated after comparison of relative expression of each gene of interest in dsRNA-injected against relative expression levels in ds*GFP*-injected female mosquitoes. The ds*ABCH2* resulted in a 75% reduction of *ABCH2* transcript level. Graphs were produced and statistically analyzed using GraphPad Prism software version 8 using Student’s t-test. For protein estimation, polypeptide extraction from legs and western blot analysis were performed as described in section “Western blot analysis”.

### Deltamethrin toxicity assays

For deltamethrin toxicity assays, 0-Day females were injected with ds*GFP* or ds*ABCH2* and then kept in cups for 72 hours as described above. After this, they were exposed to 0.016% deltamethrin (the estimated LC50 for this VK7 strain (VK7-LR) (6)), using insecticide impregnated papers in WHO tubes. The exposure lasted for 1 hour and after this the number of knocked-down mosquitoes were recorded. This was followed by 24 hour recovery in control tubes in insectary conditions. A mosquito was classified as dead or knocked down if it was immobile, unable to stand or take off. The whole procedure (injections followed by bioassays) was repeated four times (~14-16 ds*GFP* and ~18-22 ds*ABCH2* individuals x 4 replicates). In each round, ds*GFP* and ds*ABCH2* were injected concomitantly, to ensure comparable injection, insectary and bioassay conditions. Graphs were produced and statistically analyzed using GraphPad Prism software version 8 using Student’s t-test.

### ^14^ C penetration rate in ds*ABCH2* and ds*GFP* mosquitoes

Penetration rate was assessed as previously described in (41). Briefly newly-emerged ds*ABCH2*- and ds*GFP*-injected mosquitoes, three days post injection, were subjected to 4 minute contact on 0.01% ^14^ C-deltamethrin paper. Mosquitoes were collected in glass vials and three 1 minute-hexane washes were performed. After that they were homogenized in PBS. In all samples 10ml of liquid Scintillation Counting Mixture (Ultima Gold;6013326; PerkinElmer) were added and they were measured on a beta counter (LS1701; Beckman). Internal counts per minute correspond to the PBS-homogenized mosquitoes, while external counts per minute correspond to the average of the counts per minute of the three hexane washes. Penetration was calculated as the ratio of the internal to the total counts per minute in the two conditions. Two biological replicates were performed with 42 mosquitoes in each condition.

### Phylogenetic analysis

Multiple sequence alignment was performed using Mafft v7.310 (61) with default parameters. Alignments were trimmed using trimAl (62)and converted to a phylip format file using a custom Bash script. The phylogenetic tree was built under the maximum likelihood optimality criterion using IQ-TREE2 (63) with the following parameters *“-alrt 5000 -bb 5000 -m MFP”*. Tree visualization was performed using Evolview3 (64).

### Extraction of cuticular lipids, cuticular hydrocarbons (CHCs) fractionation, identification and quantitation

3 days after dsRNA nano-injections performed in newly emerged females, 95 ds*ABCH2* and 103 ds*GFP* female mosquitoes (3 replicates/ 30-35 mosquitoes per replicate) were dried at Room Temperature for about 48 h. Legs of dry mosquitoes were separated from the rest of the body and their dry weight was calculated. CHC analysis was carried out in VITAS-Analytical Services (Oslo, Norway) as described in (65). Statistics were analyzed using GraphPad Prism software, version 8 and statistical analysis carried out with Student’s t-test.

### ABCH2-overexpressing Sf9 insect cells, membrane preparations and ATPase activity estimation

ABCH2 was expressed in *Spodoptera frugiperda Sf9* insect cells using the Pfastbac1 vector, which was synthesized de novo (GenScript) and *ABCH2* ORF was subcloned in between BamHI and EcoRI restriction enzyme sites. The sequence was codon optimized for *S. frugiperda* using GenSmart codon optimization tool (GenScript). Recombinant baculoviruses encoding *ABCH2* cDNA were generated with BAC-TO-BAC Baculovirus Expression Systems (Invitrogen), following manufacturer’s instructions. 2μg of Bacmid DNA, mixed with Escort IV Transfection Reagent (Merck) were used to transfect 5×10^5^ Sf9 cells in 1ml of SF900 II SFM growth medium (ThermoFisher scientific). The complex was incubated for 45 min/RT and the lipid complexes were added dropwise to the appropriate well and were incubated at 27°C for 6 hours. After that, another 1ml of medium, supplemented with antibiotics (2x penicillin and streptomycin) and 20% FBS (ThermoFisher scientific) were added. After 24 hours DNA:lipid complexes were removed and 2ml of supplemented growth medium were added to the cells which were subsequently incubated at 27°C for 72 hours. Medium, containing the virus stock were removed and stored, while cells were also collected by pipetting in ice-cold 1 X PBS. Cell pellets (after 3000g/5min/4°C centrifugation) were resuspended in RIPA supplemented with protease inhibitors followed by a 10,000g/10 min /4°C centrifugation step. The supernatant was prepared for western blot analysis with the addition of 5x Sample Buffer (SB). After validation of recombinant protein expression of expected size in Sf9 cells and determination of viral stock titers using baculoQUANT ALL-IN-ONE (GenWay), according to manufacturer’s instructions, infection was performed. For infection 10^6^ cells/ml were seeded in T75 Flasks in a final volume of 15 ml antibiotic and FBS supplemented growth medium. After 4-5 hour incubation at 27°C, viruses were added at the desired MOI and infected cells were incubated for other 96 hours. Along with the ABCH2 infections, cells infected with an empty Bacmid and an unrelated membrane protein served as negative controls. Then the cells were harvested and proceeded to microsomal preparations. Following a centrifugation step at 2,000g/3min/4°C, cell pellets were homogenized using glass-Teflon tissue homogenizator in a mannitol-containing buffer as described in (66). Final ultracentrifugation step at 100,000g/1hour/4°C (Beckman AirFuge CLS Ultracentrifuge) resulted in pellets which were resuspended in the same buffer (66). After estimation of total protein content of the membrane preparations by Braddford protein assay using BSA to generate a linear control standard curve (0.5-20μg) the membranes were further diluted in 1-2 μg/μl to be used for downstream experiments or stored at −80°C as aliquots of 20μl each. The ATPase activity of the Sf9 membranes expressing ABCH2 or not (Empty) was estimated with a colorimetric assay by measuring inorganic Pi released from ATP hydrolysis as described by (66, 67), with some modifications. Briefly, we prepared a malachite green solution (340 mg of Malachite Green (Sigma) in 75 ml deionized water) and an ammonium molybdate solution (10.5 g ammonium molybdate in 250 ml 4N HCl). Mixing the two solutions followed by filtration through paper resulted in the malachite green stock solution that was stored at 4°C. The malachite working solution (MGWS) was freshly prepared by adding 20% Triton-X-100 in malachite green stock solution to a 0.1% final concentration. Phosphorous standard solutions incubated with malachite green working solution (50 μl of each with 0.8 μl MGWS) were used to generate linear control standard curves (0-20 nmoles Pi) (68). Pi standards were incubated in MGWS for 5 minutes and after that 100ul 37% citric acid was added in each reaction. The reactions were then incubated for 40 minutes at Room Temperature and absorbance was measured at 630nm. For estimation of Pi liberation in membrane preparations, reactions were set up using 0.5 μl of 0.1 M ATP, 2 μl of 10mg/ml BSA, 10 μg of membrane protein, 5 μl of a 10X buffer (500mM Tris/HCL Ph=8, 50mM MgCl2, 50mM KCl and 10mM DTT) and deionized water was added up to 50ul. The reactions were incubated in a water-bath at 37°C for 15 minutes to allow ATP hydrolysis to occur and afterwards they were mixed with 0.8μl freshly prepared MGWS. Then a further 5 minute incubation was performed and terminated by adding 100 μl of 37% citric acid to avoid further ATP hydrolysis. The reactions were kept at RT for 40 minutes and then absorbance measured at 630nm. In every replicate empty, ABCH2 or unrelated protein membranes were tested simultaneously. The empty Bacmid was used to remove the Sf9 non-specific ATPase background. In every ATPase experiment two control solutions were used: a reaction containing the buffer without proteins and two reactions containing no ATP, one for each membrane preparation. We used the first control as blank which was subtracted at every absorbance value and thereafter the liberated phosphate was quantified based on the Pi standard curve. As this comparison is based on total protein content estimation, whole membrane protein extracts were tested on SDS-Page and the separated polypeptides were visualized with Coomassie Blue (Figure S3C). According to the staining ABCH2 was weakly expressed in Sf9 membranes since an induced band was not apparent when expressing and control membranes were compared. For ATPase experiments with deltamethrin, the same set-up was used. Deltamethrin was resuspended in DMSO/ethanol (1:1 v/v). A working stock of 0.6 mg/ ml was used and 5μl deltamethrin was added in each reaction (3μg-120μM). Each reaction contained 10μg of total protein from empty and ABCH2 membrane isolations. Reactions containing only DMSO/ethanol were also processed. The empty/DMSO/ethanol and empty/deltamethrin Pi/μgr of total protein/minute of reaction estimated values were subtracted from the ABCH2/DMSO/ethanol and ABCH2/deltamethrin accordingly, thus removing the background and allowing estimation of ABCH2-deltamethrin stimulation over the deltamethrin controls.

### Estimation of ABCH2 content in Sf9 membranes

For the approximate estimation of ABCH2 content in Sf9 membrane preparations, western blot analysis was used, as described above. Parallel to the infection with ABCH2 bacmid, infection with a His-tagged ABCH2 bacmid was also carried out and membranes were prepared to allow quantification based on another his-tagged, purified protein of measured quantity. Total protein content of His-ABCH2 expressing membranes was quantified using Bradford protein assay and known quantities ran on the same gel with a known concentration of either a His-tagged, purified, unrelated protein or the ABCH2 membranes. Upon separation, the polypeptides were electro-transferred on two separate nitrocellulose membranes, which were blotted with anti-His or anti-ABCH2 respectively (Figure S7). Then, imageJ was used for band density analysis pairwise (His-ABCH2 and His-protein, His-ABCH2 and ABCH2). Based on the estimation described in Supplementary File 3, ABCH2 was about 11 ng per μg of total protein, which corresponds to 0.12 pmoles (based on the expected molecular weight of the transporter, 85kDa).

### Structural analysis and molecular docking simulations

The amino acid sequence of ABCH2 (Uniprot:A0A6E8VHH0) was submitted to the SWISS-MODEL in the automated protein modelling server provided by the GlaxoSmithKline center, using its standard settings (69). The same software was also used to identify the best template for the homology-based modelling (69). The human homodimeric ABCG1 transporter involved in cholesterol trafficking (42) was identified as the closest homologue. For both the ABCG1 structure and the ABCH2 predicted model, the Protein Interaction Calculator (PIC) webserver was used to predict intermolecular interactions (43). All Molecular Docking studies were performed using Autodock Smina, a fork of Autodock Vina software (includes changes from the standard Vina version 1.1.2) (46), and visualized using the PyMOL Molecular Graphics System (Version 2.0.6 Schrödinger, LLC). The structure used as receptor for the ABCG1 transporter was obtained from Protein Data Bank (PDB code: 7R8D), (70), and the 3D structures of the ligands were obtained from PubChem database (71). The nature of all protein-ligand interactions was identified using Protein-Ligand Interaction Profiler (PLIP) (71).

## Acknowledgements

We would like to thank Dr Inga Siden-Kiamos and Lefteris Spanos for providing E-cadherin antibody. We would also like to acknowledge Dr Linda Grigoraki for critical reading of the manuscript.

## Data Availability

All relevant data are within the manuscript and its Supporting information files

